# Function and Importance of Marine Bacterial Transporters of Plankton Exometabolites

**DOI:** 10.1101/2023.01.19.524783

**Authors:** William F. Schroer, Hannah E. Kepner, Mario Uchimiya, Catalina Mejia, Lidimarie Trujillo Rodriguez, Christopher R. Reisch, Mary Ann Moran

## Abstract

Metabolite exchange within marine microbial communities transfers carbon and other major elements through global cycles and forms the basis of microbial interactions. Yet lack of gene annotations and concern about the quality of existing ones remain major impediments to revealing the metabolite-microbial network. We employed an arrayed mutant library of the marine bacterium *Ruegeria pomeroyi* DSS-3 to experimentally annotate substrates of organic compound transporter systems, using mutant growth and compound drawdown analyses to link transporters to their substrates. Mutant experiments verified substrates for thirteen *R. pomeroyi* transporters. Four were previously hypothesized based on gene expression data (taurine, glucose/xylose, isethionate, and cadaverine/putrescine/spermidine); five were previously hypothesized based on homology to experimentally annotated transporters in other bacteria (citrate, glycerol, *N*-acetylglucosamine, fumarate/malate/succinate, and dimethylsulfoniopropionate); and four had no previous annotations (thymidine, carnitine, cysteate, and 3-hydroxybutyrate transporter). These bring the total number of experimentally-verified organic carbon influx transporters to 17 of 126 in the *R. pomeroyi* genome. In a longitudinal study of a coastal phytoplankton bloom, expression patterns of the experimentally annotated transporters linked them to different stages of the bloom, and also led to the hypothesis that citrate and 3-hydroxybutyrate were among the most highly available bacterial substrates. Improved functional knowledge of these gatekeepers of organic carbon uptake is facilitating better characterization of the surface ocean metabolite network.

## Introduction

The ocean microbiome plays a central role in mediating carbon and element cycles through its unique ability to process organic carbon dissolved in seawater (1-3). Ultimately, marine bacteria take up and assimilate as much as half of marine net primary production (NPP) in the form of exometabolites derived from excretion and death of phytoplankton and other microbes (3, 4). Given that current and future controls over this globally important carbon flux are poorly understood, identification of the metabolites produced and consumed by ocean microbes is critically needed (5).

One approach to unraveling marine metabolite flux is through the application of transcriptomic and proteomic tools by which dynamics of the chemical environment can be gleaned from changes in the expression of microbial genes. Such approaches are easy to scale with advancements in sequencing and data sharing (6, 7) and have successfully addressed metabolite dynamics in various microbial systems such as model communities (8), phytoplankton blooms (9, 10), oligotrophic ocean regions (11, 12), and global-scale ocean surveys (13-15). Transporter genes in particular are of value in such approaches because they are a cell’s interface with its environment and their expression can reveal the identity of available metabolites (16). A key limitation to their use, however is a dependence on accurate gene annotation to identify protein function. For most microbial transporters, the substrate is still unknown. Others are annotated computationally based on homology (17-19), yet this is error prone when relationships to experimentally annotated genes is distant (20). Indeed, transporters have a lower rate of successful annotation based on homology than catabolic enzymes (19).

Experimental confirmation of gene annotation is the gold standard, but is both time and resource intensive. Moreover, it is largely limited to cultured species for which genetic systems are available, leaving out much of the diversity represented in environmental bacteria. An alternate approach uses pooled transposon mutants whose fitness under defined selection pressure provides a hypothesis of gene function (21-24). This method requires only a minimal genetic system to introduce small DNA fragments (transposons) and a protein that catalyzes genomic insertion (transposase) into bacterial cells. A recent high-throughput advancement of this method, termed BarSeq (25, 26), uses unique barcodes that link each transposon insertion site to the specific gene it disrupts, thereby allowing mutant pools to be analyzed for fitness through cost-effective amplicon sequencing. A wide taxonomic range of bacteria have been shown to be amenable to BarSeq library construction, resulting in hypothesis generation for gene functions that include stress response, metabolism, phage resistance, and transport (25, 27-29). Hypotheses can be confirmed experimentally if targeted single-gene mutants are subsequently constructed, as for those predicting membrane proteins (28) and catabolic enzymes (30).

For a small number of well-studied model bacterial species, genome-wide arrayed mutant libraries have been constructed through painstaking targeted gene deletions to produce libraries of single-gene knockouts across the genome. Excellent tools for gene annotation, these arrayed libraries are currently available for well-established model bacteria, such as *Escherichia coli* (31), *Acinetobacter baylyi* (32), *Bacillus subtilis* (33, 34), and *Salmonella enterica* (35). Pooled transposon mutant libraries have been used successfully as the starting material for such arrayed libraries (24, 36, 37), but require individual sequencing of tens of thousands of colonies to determine transposon insertion location.

Recently, a modification of the BarSeq approach was used to create an inexpensive arrayed mutant library of the marine model bacterium *Ruegeria pomeroyi* DSS-3 (38). The method took advantage of the ease of insertion site identification in BarSeq libraries in combination with 384-well plate location barcodes to generate an arrayed library of single-gene knockout mutants for *R. pomeroyi*, similar to the approach used recently for the anaerobic gut microbe *Bacteroides thetaiotaomicron* (39). *R. pomeroyi* is known for its ecological association with marine phytoplankton and ability to grow on plankton-derived metabolites (16, 40, 41), but to this point substrates of only four of the 126 putative organic compound influx transporters have been experimentally verified via gene knockout mutants: choline (42), dihydroxypropanesulfonate (DHPS) (41, 43), ectoine (44), and trimethylamine N-oxide (13). Here we leverage a set of 156 influx transporter mutants from the arrayed *R. pomeroyi* BarSeq (arrayed-BarSeq) library in high-throughput screens against 63 possible substrates to increase knowledge of transporter function. Resulting gene annotations were then applied to a set of *R. pomeroyi* transcriptomes sampled after introduction to a bloom Monterey Bay, CA, USA (Nowinski and Moran, 2021). The 13 newly verified transporter annotations provided insights into the metabolites serving roles as substrates to heterotrophic bacteria during a coastal bloom.

## Methods

### BarSeq library generation and mapping

Full methods for generating and arraying the *R. pomeroyi* BarSeq mutant library are provided in Mejia *et al*. (38). Briefly, a pool of randomly barcoded transposon mutants was constructed according to Wetmore *et al*. (26) by conjugating *R. pomeroyi* DSS-3 with *E*.*coli* WM3064 containing the transposome pKMW7 Tn5 library (strain APA766). The insertion sites were subsequently linked to the unique barcodes by sequencing through the barcoded transposons into the disrupted genes.

To construct the arrayed libraries, individual mutants were isolated on ½ YTSS solid medium amended with 100 µg ml^-1^ kanamycin. Colonies were picked after 2 d (Qpix2 automated colony picker; Molecular Devices, San Jose, CA) and arrayed into 384 well plates containing 80 µl of liquid ½ YTSS + kanamycin medium. Plates were incubated at 30°C for 2-5 d until visible growth appeared and then replicated.

Glycerol was added to a final concentration of 20% and plates were frozen at −80°C. To identify the mutant situated in each well, a set of 16 forward and 24 reverse location primers were synthesized with unique 8 bp barcodes. These were used combinatorially in 384 unique pairs for PCR amplification of the 20 bp BarSeq barcodes linked to the 8 bp location primers, mapping mutants to their well location. In total 27,488 unique mutants were arrayed in 384-well plates, covering 3,292 protein encoding genes. Mutants for 156 putative organic compound influx transporter genes were re-arrayed into two 96 well plates for subsequent screening (Table S1).

### Growth Screen

Mutant cultures were pre-grown overnight in ½ YTSS medium with 50 μg ml^-1^ kanamycin. Screens were performed in L1 minimal medium (45) modified to a salinity of 20 and amended with ammonium (3 mM) and kanamycin (50 μg ml^-1^). For the initial screen, overnight cultures of individual mutants (2 μl) were inoculated into 198 μl of modified L1 with a single substrate as the sole carbon source at 8 mM carbon. Plates were incubated at 25°C with shaking, and optical density (OD_600_) was read at intervals of 6-12 h until cultures entered stationary phase at ∼24-48 h. Mutants exhibiting phenotypes in the initial screen were moved to the targeted screen in which 4 replicate 200 µl mutant cultures were prepared by inoculating 2 μl of washed (3x) overnight culture into 96 well plates containing 198 μl modified L1 medium and a substrate at 8 mM carbon. As a positive control, four wells with the same medium were inoculated with washed overnight cultures of the pooled-BarSeq library, used as a proxy for wild-type *R. pomeroyi* growth but harboring a transposon/kanamycin resistance gene insertion. Cultures were grown at 25°C in a Synergy H1 plate reader (BioTek, Winooski, VT, USA) shaking at 425 rpm for 68-72 h. OD_600_ readings were corrected to a pathlength of 1 cm assuming a volume of 200 µl.

Mutant defect was identified by comparison to the OD_600_ achieved by the pooled-BarSeq library (n=4; ANOVA and TukeyHSD; p ≤ 0.05) (Table 1). Mutants with significantly lower OD_600_ on multiple substrates were regrown on rich medium to check for viability, and removed from further consideration if they broadly demonstrated poor growth; two mutants were removed after this viability check (SPO0050 and SPO2952).

**Table 1.** Transporter identification based on growth and metabolite drawdown screens. ΔOD, percent decrease in optical density of the isolated mutant relative to pooled-BarSeq library with associated 95% confidence interval and p value (n=4, ANOVA with TukeyHSD). ΔDrawdown, percent decrease in drawdown by the isolated mutant relative to the pooled-BarSeq library with associated 95% confidence interval and p value (n=3, ANOVA with TukeyHSD). Prediction, previous annotation status of the transporter as follows: novel = annotation was not known or hypothesized; homology = annotation was hypothesized based on sequence similarity; expression = annotation was hypothesized based on gene expression data; control = annotation known based on previous *R. pomeroyi* knockout mutant. GlcNAc, *N*-acetylglucosamine, N.S., not significant (p≥0.05).

| Transporter | Mutant | Substrate | Prediction | $\Delta OD$ (%) | 95% CI (% +/-) | p adj | ΔDrawdown (%) | 95% CI (% +/-) | p adj |
| --- | --- | --- | --- | --- | --- | --- | --- | --- | --- |
| *tctABC* | ΔSPO0184 | Citrate | homology | 99.1 | 12.4 | $1.1 \times 10^{-6}$ | 99.3 | 3.6 | $5.4 \times 10^{-12}$ |
| *nupABC* | ΔSPO0378 | Thymidine | novel | 81.1 | 23.5 | $1.3 \times 10^{-5}$ | 63.6 | 32.5 | $2.3 \times 10^{-3}$ |
| | ΔSPO0379 | Thymidine | novel | 81.5 | 23.5 | $1.3 \times 10^{-5}$ | 83.0 | 32.5 | $5.6 \times 10^{-4}$ |
| *hpsKLM* | ΔSPO0591 | DHPS | control | 78.6 | 30.7 | $7.7 \times 10^{-4}$ | 64.8 | 19.0 | $1.1 \times 10^{-4}$ |
| *glpVSTPQ* | ΔSPO0608 | Glycerol | homology | 79.6 | 13.0 | $5.3 \times 10^{-6}$ | 78.8 | 2.8 | $2.1 \times 10^{-12}$ |
| *tauABC* | ΔSPO0674 | Taurine | expression | 91.8 | 20.6 | $1.5 \times 10^{-6}$ | -3.1 | 34.2 | N.S. |
| | ΔSPO0676 | Taurine | expression | 92.8 | 20.6 | $1.4 \times 10^{-6}$ | 49.1 | 34.2 | $1.1 \times 10^{-2}$ |
| *xylFGH* | ΔSPO0863 | Glucose | expression | 92.1 | 9.7 | $4.9 \times 10^{-7}$ | 85.6 | 30.9 | $1.5 \times 10^{-3}$ |
| | ΔSPO0863 | Xylose | expression | 96.6 | 5.6 | $1.1 \times 10^{-8}$ | 52.8 | 45.5 | $3.2 \times 10^{-2}$ |
| *betT* | ΔSPO1087 | Choline | control | 100.5 | 8.9 | $2.3 \times 10^{-7}$ | 94.1 | 10.8 | $2.2 \times 10^{-5}$ |
| *uehABC* | ΔSPO1147 | Ectoine | control | 86.7 | 6.1 | $5.5 \times 10^{-8}$ | 62.1 | 3.8 | $9.2 \times 10^{-3}$ |
| *nagTUVW* | ΔSPO1839 | GlcNAc | homology | 78.4 | 7.1 | $2.6 \times 10^{-7}$ | 92.7 | 5.0 | $5.7 \times 10^{-8}$ |
| *iseKLM* | ΔSPO2357 | Isethionate | expression | 101.1 | 10.2 | $1.8 \times 10^{-9}$ | 96.0 | 41.9 | $1.0 \times 10^{-3}$ |
| | ΔSPO2358 | Isethionate | expression | 104.2 | 10.2 | $1.6 \times 10^{-9}$ | 69.8 | 41.9 | $5.3 \times 10^{-3}$ |
| *hbtABC* | ΔSPO2573 | 3-OH butyrate | novel | 92.7 | 3.9 | $5.4 \times 10^{-10}$ | 79.8 | 11.7 | $5.3 \times 10^{-5}$ |
| *dctMPQ* | ΔSPO2626 | Fumarate | homology | 99.1 | 3.3 | $2.7 \times 10^{-14}$ | 98.0 | 7.9 | $1.6 \times 10^{-9}$ |
| | ΔSPO2626 | Malate | homology | 48.6 | 25.3 | $4.9 \times 10^{-4}$ | 58.1 | 13.6 | $3.8 \times 10^{-6}$ |
| | ΔSPO2626 | Succinate | homology | 92.6 | 11.0 | $3.6 \times 10^{-11}$ | 64.9 | 5.3 | $1.9 \times 10^{-9}$ |
| | ΔSPO2628 | Fumarate | homology | 96.7 | 3.3 | $2.7 \times 10^{-14}$ | 95.0 | 7.9 | $2.1 \times 10^{-9}$ |
| | ΔSPO2628 | Malate | homology | 11.2 | 25.3 | N.S. | 47.7 | 13.6 | $1.7 \times 10^{-5}$ |
| | ΔSPO2628 | Succinate | homology | 75.2 | 11.0 | $6.9 \times 10^{-10}$ | 50.8 | 5.3 | $9.6 \times 10^{-9}$ |
| | ΔSPO2630 | Fumarate | homology | 96.9 | 3.3 | $2.7 \times 10^{-14}$ | 94.9 | 7.9 | $2.1 \times 10^{-9}$ |
| | ΔSPO2630 | Malate | homology | 12.7 | 25.3 | N.S. | 62.7 | 13.6 | $2.1 \times 10^{-6}$ |
| | ΔSPO2630 | Succinate | homology | 81.8 | 11.0 | $2.3 \times 10^{-10}$ | 58.0 | 5.3 | $4.4 \times 10^{-9}$ |
| *cntTUVWX* | ΔSPO2995 | Carnitine | novel | 48.1 | 25.8 | $1.4 \times 10^{-3}$ | 69.2 | 36.7 | $2.8 \times 10^{-3}$ |
| | ΔSPO2996 | Carnitine | novel | 61.1 | 25.8 | $2.5 \times 10^{-4}$ | 85.7 | 36.7 | $9.1 \times 10^{-4}$ |
| *cuyTUVW* | ΔSPO3040 | Cysteate | novel | 50.3 | 15.3 | $1.9 \times 10^{-5}$ | 38.9 | 30.7 | $1.9 \times 10^{-2}$ |
| | ΔSPO3041 | Cysteate | novel | 81.3 | 15.3 | $3.4 \times 10^{-9}$ | 40.0 | 30.7 | $1.7 \times 10^{-2}$ |
| *dmdT* | ΔSPO3186 | DMSP | homology | 48.9 | 13.4 | $1.1 \times 10^{-4}$ | 33.6 | 2.4 | $1.6 \times 10^{-6}$ |
| *potFGHI* | ΔSPO3469 | Cadaverine | expression | 62.3 | 7.4 | $8.3 \times 10^{-7}$ | 61.2 | 23.5 | $1.9 \times 10^{-3}$ |
| | ΔSPO3469 | Putrescine | expression | 68.8 | 10.3 | $3.2 \times 10^{-6}$ | 65.1 | 12.8 | $1.5 \times 10^{-4}$ |
| | ΔSPO3469 | Spermidine | expression | 63.6 | 16.8 | $8.9 \times 10^{-5}$ | 75.2 | 5.8 | $2.4 \times 10^{-6}$ |

### Metabolite drawdown screen

For each mutant-substrate pair identified from the growth screens, 3 replicate 220 µl cultures were prepared in 96 well plates by inoculating 3 μl of washed (3x) overnight mutant cultures into minimal medium containing the candidate substrate at 8 mM carbon. Cultures were grown shaking at 25°C for 24 h or 36 h, depending on the growth rate supported by the carbon source. At termination, 200 µl of medium were collected and centrifuged at 3,700 rpm for 10 min, and the supernatant was stored at - 80°C. Metabolite analysis was performed using a Bruker Avance lll 600 MHz spectrometer (Bruker, Billerica, MA, USA) equipped with a 5-mm TCI cryoprobe. Samples were prepared with addition of a deuterated phosphate buffer (30 mmol L^-1^, pH 7.4) and the internal standard 2,2-dimethyl-2-silapentane-5-sulfonate-d_6_ (DSS, 1 mmol L^-1^) (10:1 (vol : vol)) and transferred to 3 mm NMR tubes (Bruker). Data were acquired by a one dimensional ^1^H experiment with water suppression (noesypr1d, Bruker) at 298K using TopSpin 3.6.4 (Bruker). For glycerol, a ^1^H *J*-resolved experiment (jresgpprqf) was used to avoid overlapping background peaks. Spectra were processed using NMRPipe on NMRbox (46, 47), and the processed data were analyzed using Metabolomics Toolbox (https://github.com/artedison/Edison_Lab_Shared_Metabolomics_UGA) and MATLAB R2022a (MathWorks). For quantification of metabolites, spectra were normalized to DSS and peak area for representative peaks was calculated. TopSpin experiment settings, NMRpipe spectra processing parameters, and MATLAB data analysis scripts are available in Metabolomics Workbench (see Data Availablity).

### Pooled-BarSeq experiment

Minimal medium was prepared for 23 substrates (Fig. 3) at 8 mM carbon in a 96 well plate (n=4). Each well was inoculated with 20 μl of washed (3x) overnight culture of the *R. pomeroyi* pooled-BarSeq library. After growth with shaking at 25°C for 72 h, cultures were serially transferred into fresh media four additional times and then transferred to 1.5 ml tubes, pelleted by centrifugation at 8,000 x g for 3 min, and stored at −80°C until further processing. Genomic DNA was extracted from the cell pellets using the DNEasy blood and tissue kit (Qiagen, Hilden, Germany). PCR amplification of BarSeq barcodes was performed using primers modified from Wetmore *et al*. (26) with PhusionHF master mix (Fisher, Pittsburg, PA). An aliquot of 8 ng of product from each sample was pooled, purified using HiPrep beads (MagBio, Gaithersburg, MD, USA), and sequenced on a NextSeq SE150 Mid Output flow cell (SE150) at the Georgia Genomics and Bioinformatics Core Facility (Athens, Georgia, USA). Sequence data were processed according to Wetmore *et al*. (26). Following quality control, an average of 35,090 unique barcodes mapped to insertions that fell within the interior 10 to 90% of *R. pomeroyi* coding sequences. In total, 55 million reads were mapped to insertions in 3,570 genes (out of 4,469 protein-encoding genes in the *R. pomeroyi* genome) with a median of 404,513 mapped reads per sample. Reads mapping to different insertion sites within the same coding sequence were pooled for subsequent analyses. Mutant enrichment or depletion relative to initial abundance was used as a proxy for fitness, calculated as the mean fold-difference between abundance in a given treatment compared to abundance in all other treatments.

### Transporter expression during a Monterey Bay bloom

Processed *R. pomeroyi* transcriptome data (transcripts per million and Z-scores), metadata, and complete experimental methods are available elsewhere (9). Briefly, on 14 days over 5 weeks, *R. pomeroyi* cells were added to 350 ml of unfiltered surface water (n=3). *R. pomeroyi* was inoculated at cell numbers equivalent to that of natural heterotrophic bacteria. Subsequent sequencing analysis indicated that *R. pomeroyi* transcripts averaged 38% of the bacterial reads in the metatranscriptome datasets (48). Cells were collected by filtration after 90 min and processed for RNAseq analysis.

### Homologs in the Roseobacter group

Roseobacter strains with complete genomes available through RefSeq were selected based on Simon *et al*. (49). Phylogenic analysis of the 14 selected strains was carried with a set of 117 single copy genes using GToTree v1.6.37 (50). *R. pomeroyi* transporter genes with homologs in the other strains were identified by BLASTp using Diamond v2.0.14.152 (51), threshold: E ≤ 10^−5^ and identity ≥ 70%. Data analysis and figure generation was performed using R v3.6.1. Manual checks of gene neighborhoods were performed when BLASTp results showed that multicomponent transporters were missing one or more component gene.

## Results and Discussion

From a pooled-BarSeq transposon mutant library of *R. pomeroyi* prepared according to Wetmore *et al*. (26), 48,000 colonies were individually arrayed into 384 well plates (Fig. 1). The gene disrupted in each arrayed mutant was determined by sequencing the transposon barcode in conjunction with indexed primers that indicated plate column and row (38), creating a library that covers 3,292 of the 4,469 protein-encoded genes in the *R. pomeroyi* genome (73%). From the genome annotations (52, 53) we identified 156 mutants that were predicted to encode for 104 organic compound influx transporter proteins (Table S1). These were re-arrayed into multi-well plates to facilitate functional screens on 63 compounds known to be produced by marine phytoplankton (54).

**Fig. 1.**
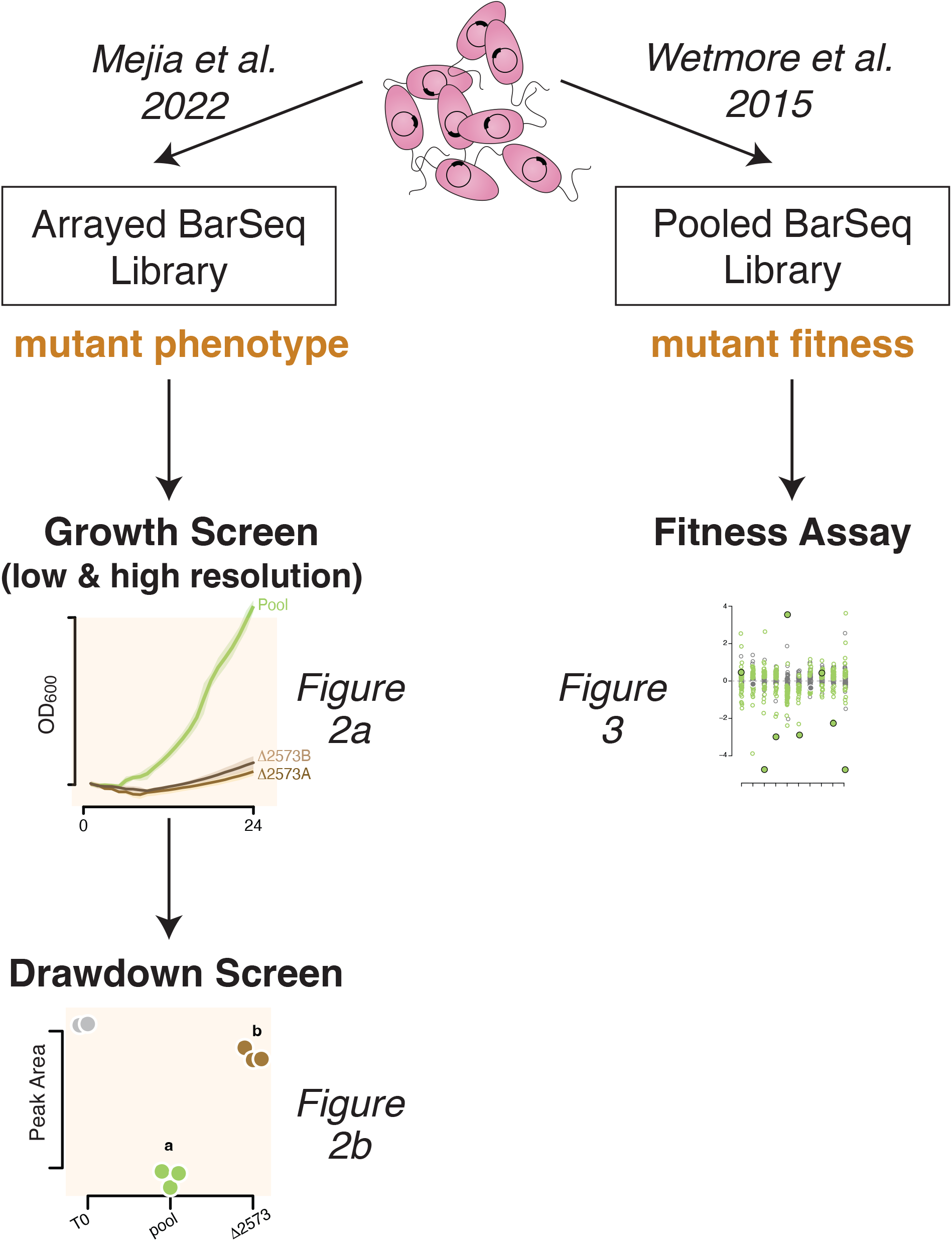
Experimental flow chart for the *R. pomeroyi* BarSeq library. Right path: Pooled mutant populations (pooled-BarSeq library) are used for gene fitness assays. Left path: Individual transporter mutants from the library (arrayed-BarSeq library) are used to screen for growth and metabolite drawdown.

### Growth Screens

Initial screens of the 156 mutants identified candidate substrates of transporter genes based on OD_600_ deficits after 24-72 h) (n=∼2). These mutants were transferred to a second round of screening in which each candidate substrate/mutant pair was monitored for growth with hourly OD readings and higher replication (n=4). A positive control treatment consisting of the full pooled-BarSeq library approximated wild-type growth (Fig. 2A, Fig. S1). We used mutants of three previously confirmed transporters as positive controls for the screening protocols; these were the *tctABC* for choline uptake (42), *hpsKLM* for DHPS uptake (41, 43), and *uehABC* for ectoine (44) (Table 1) (Fig. S1). Mutants that exhibited growth deficits on more than one metabolite were not considered further unless the metabolites had high structural similarity.

**Fig. 2.**
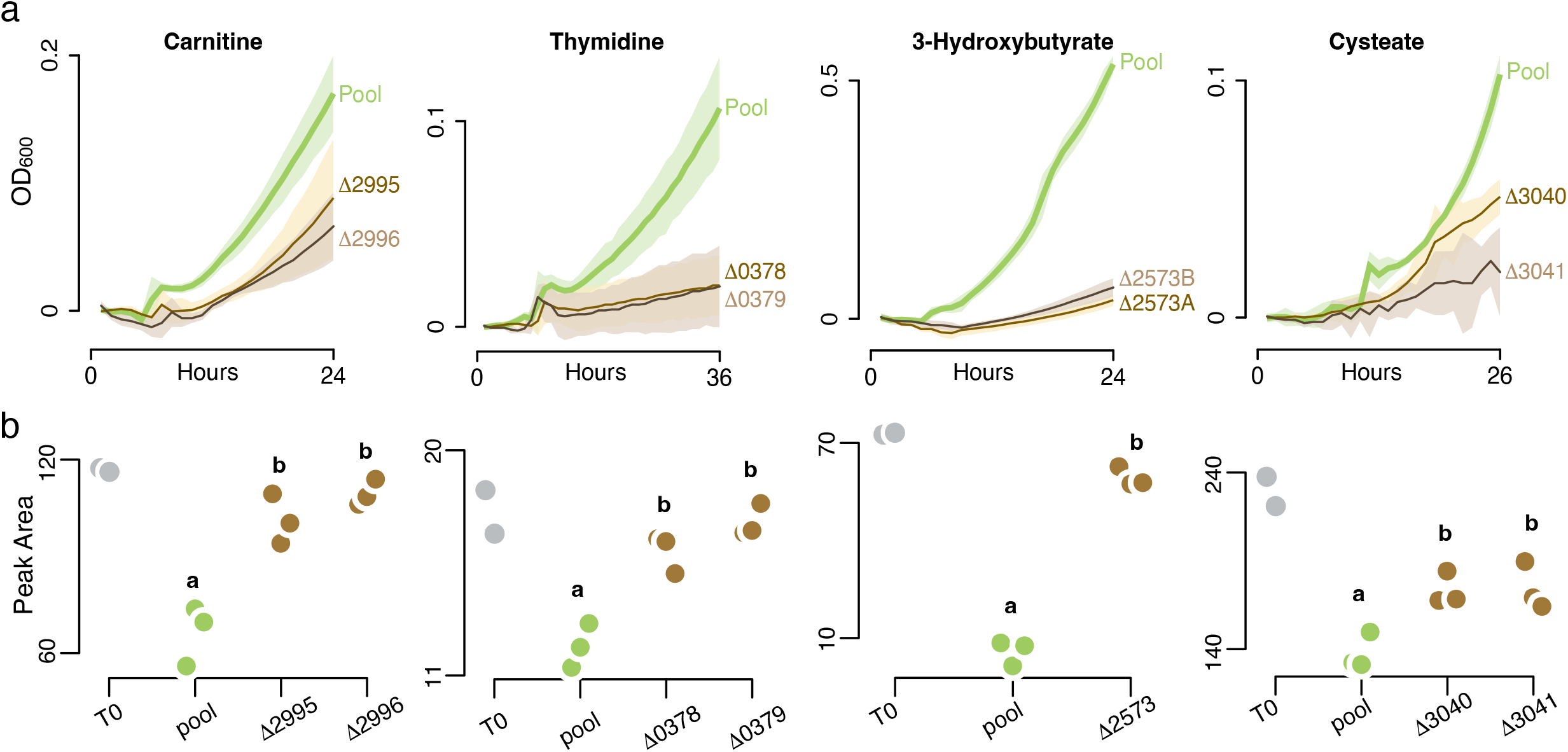
Growth and substrate draw-down results for the four novel transporter annotations. a) Growth of transporter mutants compared to growth of the pooled-BarSeq library (an analog for wild-type growth but carrying transposon and resistance gene insertions) on selected marine plankton metabolites. Shaded regions indicate 95% confidence intervals (n=4). Numbers refer to *Ruegeria pomeroy*i DSS-3 locus tags (Table 1). B) Substrate concentrations (^1^H-NMR peak area) after growth of mutants (brown symbols, n=3) or the pooled-BarSeq library (green symbols, n=3), and at inoculation (gray symbols, n=2). Letters that differ indicate that peak area for the isolated mutant(s) was significantly different than for the pooled-BarSeq library (ANOVA, n=3, p ≤ 0.05), with a TukeyHSD test carried out when multiple mutants for the same substrate were tested (p ≤ 0.05). For full results, see Table 1.

The growth-based screening process resulted in substrate predictions for 13 *R. pomeroyi* transporters (Fig. 2A, Fig. S1, Table 1). Four of these were consistent with target metabolites hypothesized based on previous gene expression data: *xylFGH* (glucose/xylose) (55), *iseKLM* (isethionate) (56), *potFGHI* (polyamines: cadaverine, spermidine, and/or putrescine)(57), and *tauABC* (taurine) (58). Four were consistent with target metabolites hypothesized based on *in silico* analysis by the GapMind tool for carbon sources (19): *tctABC* (citrate), *dctMPQ* (the C4 organic acids succinate, fumarate, and malate), *nagTUVW* (*N*-acetylglucosamine), and *glpVSTPQ* (glycerol). One was consistent with a target metabolite based on homology to an experimentally verified transporter in the closely related species *Roseovarius nubinhibens* (59): *dmdT* (dimethylsulfoniopropionate (DMSP) (Table 1). Four were novel annotations with no previous substrate predictions: *cntTUVWX* (carnitine), *cuyTUVW* (cysteate), *hbtABC* (3-hydroxybutyrate), and *nupABC* (thymidine) (Table 1). All hypothesized substrates were identified in previous studies as endometabolites in cultured phytoplankton or natural plankton communities (40, 60), or as exometabolites in phytoplankton cultures or seawater (61, 62).

### Metabolite Drawdown Screens

Substrate identifications emerging from the growth screens were further tested in metabolite drawdown experiments. Similar to the design of the growth screens, isolated mutants were inoculated into minimal medium with a single substrate as the sole carbon source (n=3), alongside positive control treatments inoculated with the pooled-BarSeq library as an analog for wild type. Spent media samples were collected at 24 h or, for substrates that supported slower growth, at 36-48 h (Fig. 2B, Fig S2).

Substrate concentration was measured by ^1^H-NMR and a mutant drawdown defect was defined as significantly higher substrate concentration in the mutant cultures compared to the pooled-BarSeq library (ANOVA and TukeyHSD, p ≤ 0.05, Table 1). All transporter annotations that had emerged from the growth screens were subsequently upheld in these draw-down screens (Fig. 2B, Fig S2), consistent with gene disruption reducing or eliminating substrate uptake (Fig. 2A, Fig S1).

Some transporter mutants, such as *betT*, were completely unable to grow on or draw down the substrate (Figs. S1, S2). This is the expected pattern if the disrupted transporter is the only system for uptake by *R. pomeroyi*. Alternatively, some of the transporter mutants, such as *dmdT*, were capable of partial growth and draw-down, but significantly less than the mutant pool (Figs. 2, S1, S2). This pattern suggests that more than one transporter in the *R. pomeroyi* genome can take up the compound. For example, *dmdT* belongs to the BCCT-type family whose members frequently have low substrate affinity (63), suggesting that a second, high-affinity transporter may be used when substrates become depleted; previous studies have similarly suggested that *R. pomeroyi* may have more than one DMSP transporter (59, 64). In a mixed result, complete loss of growth and draw-down for fumarate yet partial losses for succinate and malate suggests that *dctMPQ* is the only transporter system in the *R. pomeroyi* genome for fumarate, but the other C4 organic acids likely have a second transporter (Fig. 2B, Fig S2, Table 1).

### Comparison to Pooled-BarSeq Libraries

Another approach to identify substrates of bacterial transporters is to place a pooled-BarSeq library under selection on a single carbon source (25). In this case, transporter mutants that exhibit poor growth are identified as candidate uptake systems. We asked whether the pooled-BarSeq approach would have been sufficient to recognize the *R. pomeroyi* transporters identified here, saving the effort of arraying the BarSeq library while also providing additional information on catabolic and regulatory genes that may support metabolite utilization.

Mutant abundance was calculated for members of the pooled-BarSeq library following selection for growth on ten substrates used in the growth screens (Fig. 3). Selection occurred over four growth dilution cycles of 72 h each. Amplicon sequencing of the pooled library at the beginning and end of selection (26) was used to calculate relative growth rates for each mutant in the pool as a proxy for fitness. For five substrates, the pooled BarSeq results agreed with results from the arrayed mutant screens, identifying the same transporter systems for DHPS, ectoine, glucose, 3-hydroxybutyrate, and spermidine (n=4; T test, p ≤ 0.05) (Fig. 3). For five other substrates, the known transporter mutant was either not significantly depleted from the mutant pool or significantly enriched, and thus transporters were not correctly identified for cysteate, DMSP, *N*-acetylglucosamine, xylose, and malate. In a counterintuitive finding, the *N*-acetylglucosamine transporter mutant *nagTUVW* was the most enriched population in the pool, indicating a fitness gain for cells unable to take up the only provided substrate. We hypothesize that this was due to cross-feeding of an *N*-acetylglucosamine degradation product, released by the other mutants whose knockouts are in unrelated genes. While these results demonstrate that pooled-BarSeq mutant libraries are excellent tools for low-cost, high-throughput hypothesis generation, predicted transporter annotations nonetheless require experimental follow-up (28, 30).

### Transporter Expression in a Coastal Phytoplankton Bloom

We used an *R. pomeroyi* gene expression dataset from a natural phytoplankton bloom in Fall 2016 in Monterey Bay, CA, USA (48) to assess the ecological relevance of the verified transporters. On 14 dates over 5 weeks during the decline of a bloom dominated by the dinoflagellate *Akashiwo sanguinea, R. pomeroyi* cells were introduced into the natural community for 90 min (9). Metatranscriptomic data from each sample were subsequently mapped to the *R. pomeroyi* genome to identify environmental conditions eliciting transcriptional responses. We reanalyzed this dataset in light of the new information on transporter function, with the goal of generating insights into bloom-associated metabolites supporting heterotrophic bacterial growth.

To first evaluate the internal consistency of the expression data, pairwise correlation coefficients were calculated across the sample dates for the individual components of the 14 multi-component transporters. Nine systems had within-transporter correlation coefficients above 0.84 (Pearson correlation, p ≤ 0.05), confirming coherence in the expression patterns for genes in the same transporter system (Fig. 4a). The remaining four had within-transporter correlation coefficients ranging from 0.10 to 0.60; three of these, however, had particularly low expression in Monterey Bay (Fig. 4b) that may have affected the accuracy of expression calculations.

**Fig. 3.**
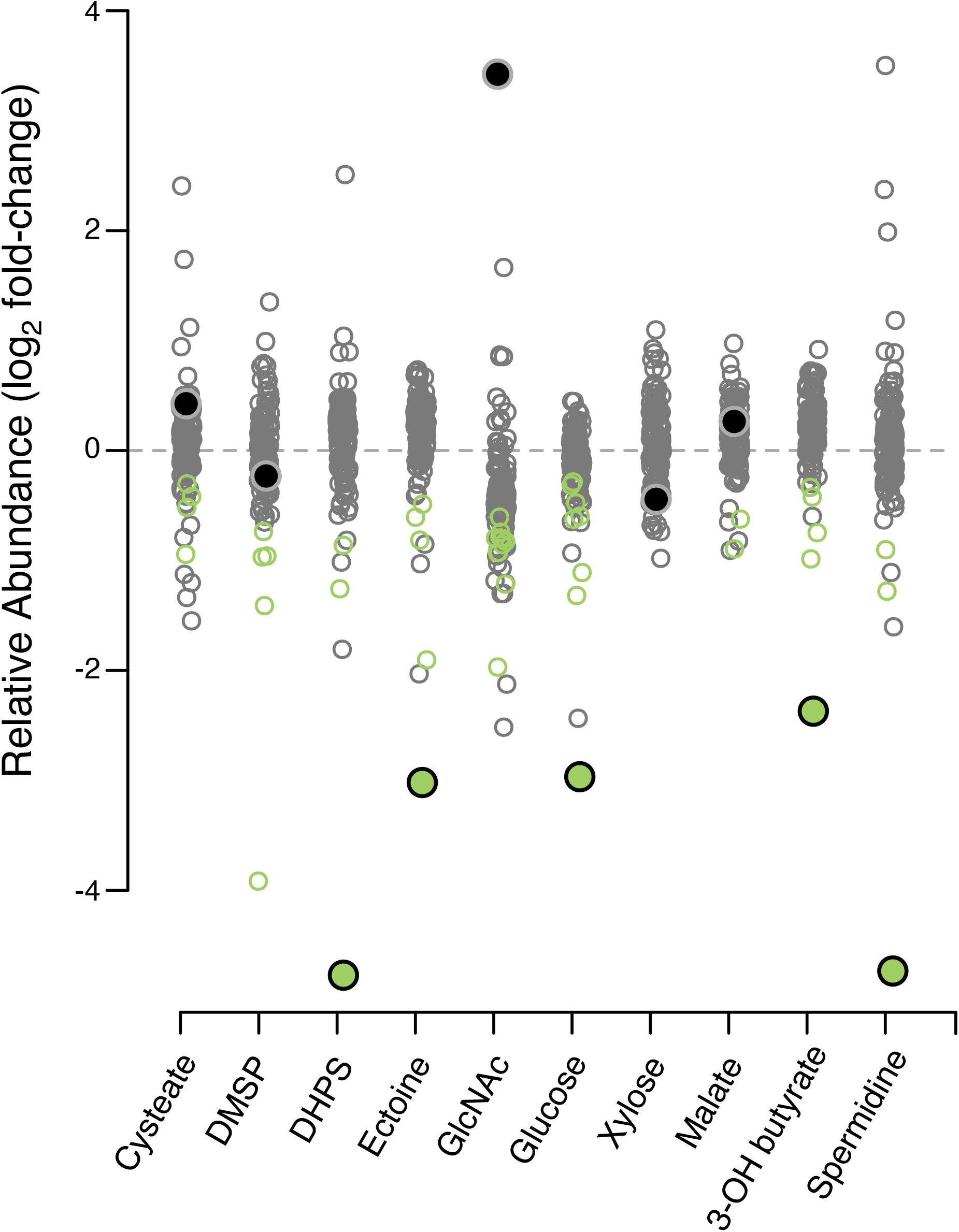
Relative abundance of *Ruegeria pomeroyi* DSS-3 transporter mutants following selection of the pooled-BarSeq library for growth on 10 metabolites. Green symbols indicate significant mutant depletion (T-test, n=4, Benjamini-Hochberg adjusted p ≤ 0.05) and gray symbols indicate non-significant changes. The larger filled symbols indicate the identified transporter for that metabolite as determined from growth and draw-down assays of individual mutants, and is colored green if it was correctly identified, and colored gray if not. Mutant enrichment/depletion for multi-gene transporter systems is plotted as the average of all components.

**Fig. 4.**
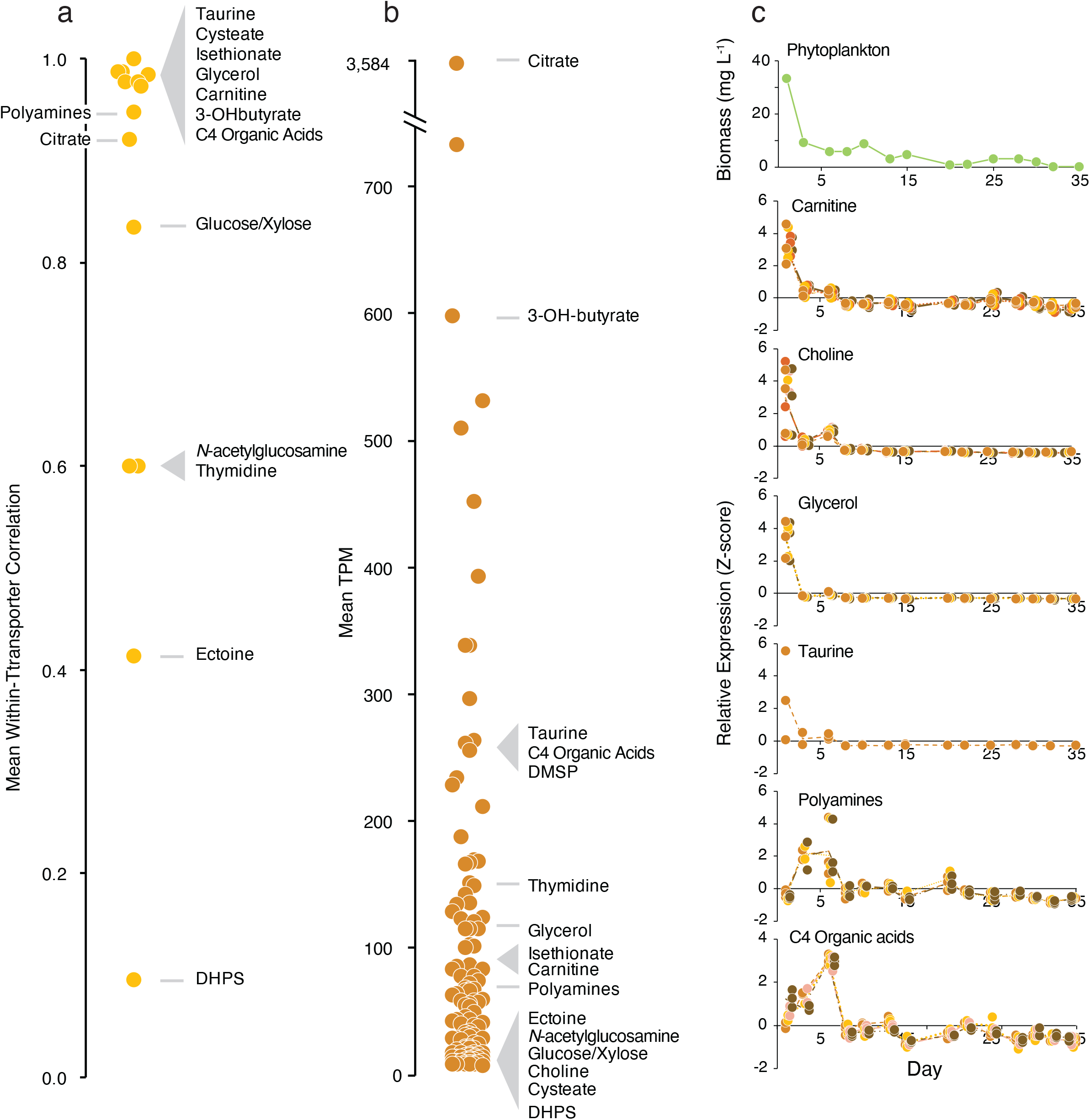
a) Mean correlation coefficients of relative expression levels of genes in multi-component transporter systems in Monterey Bay, in Fall, 2016. b) Mean relative expression levels for the 126 *R. pomeroyi* transporter systems when introduced into Monterey Bay seawater, averaged across 14 dates. c) Relative expression of selected *R. pomeroy*i DSS-3 transporters in Monterey Bay seawater on each of 14 dates over 35 days, normalized as Z-scores. Transporters have 1-5 component genes (each colored in different shades of brown) and each component gene has three replicates plotted individually. Lines connect the component mean expression through time. Total phytoplankton biomass (μg C L^-1^) during the 5-week sampling period is also shown.

Expression patterns of the carnitine, choline, taurine, and glycerol transporters were positively related to phytoplankton biomass through the bloom (Pearson correlation, p ≤ 0.05) (Fig. 4c), and we hypothesize that these compounds are consistent members of the exometabolite pool in dinoflagellate-dominated blooms. Expression of the C4 organic acid and polyamine transporters had peak expression coinciding with the largest drop in phytoplankton biomass (Fig. 4c), and we hypothesize that these compounds are released from senescing or dead phytoplankton. Transcripts from *R. pomeroyi*’s 126 transporter systems were ranked by their abundance in the transcriptomes [mean transcripts per million (TPM), averaged across components for multi-gene transporters]. If heterotrophic bacterial transporter expression is regulated by substrate detection (admittedly an oversimplification (65)), citrate, 3-hydroxybutyrate, taurine, and DMSP, were among the most important sources of organic carbon to *R. pomeroyi* in this bloom (expression ranked in the top 25% of transporters). Conversely, DHPS and cysteate were among the least important (ranked in the bottom 25%) (Fig. 4b). 3-hydroxybutyrate was of particular interest for two reasons. First, transport systems for this substrate are poorly understood (66) with *hbtABC* representing the only confirmed identification of a bacterial transporter for this metabolite. Second, *hbtABC* was the third most-highly expressed *R. pomeroyi* transporter in Monterey

Bay, indicative of an unrecognized role as an important bacterial carbon source. The most highly expressed of all the *R. pomeroyi* transporters, however, was citrate, averaging almost 5-fold higher relative transcript abundance than the next highest (Fig. 4b).

### Homologous Transporters in the Roseobacter group

*R. pomeroyi* and its relatives in the Roseobacter group are recognized for high abundance in many coastal marine environments (67, 68). The cultured members of this group typically have large, well-regulated genomes capable of diverse metabolisms (69) and are often associated with phytoplankton blooms (68, 70, 71). To determine the distribution of the 17 verified transporters in Roseobacter genomes, 13 other strains with closed genomes and representing a broad sampling of the group’s phylogenetic diversity (49) were selected for analysis. Transporters for *N-*acetylglucosamine, C4 organic acids, polyamines, and carnitine are present only in close relatives of *R. pomeroyi*, consistent with vertical transmission (Fig. 5). Transporters for the organic sulfur compounds DHPS, taurine, and isethionate are common in deeply branching strains but retained in few of the more recently branching lineages. The transporters for cysteate and ectoine are unique or nearly so to *R. pomeroyi*, suggestive of specialized niche dimensions. Finally, transporters for thymidine, citrate, glycerol, and 3-hydroxybutyrate are well conserved throughout Roseobacter genomes (Fig. 5), indicating broad importance of these substrates to the ecology of this group. Patchy distribution of transporter orthologs relative to the group’s phylogeny has been reported previously (72).

**Fig. 5.**
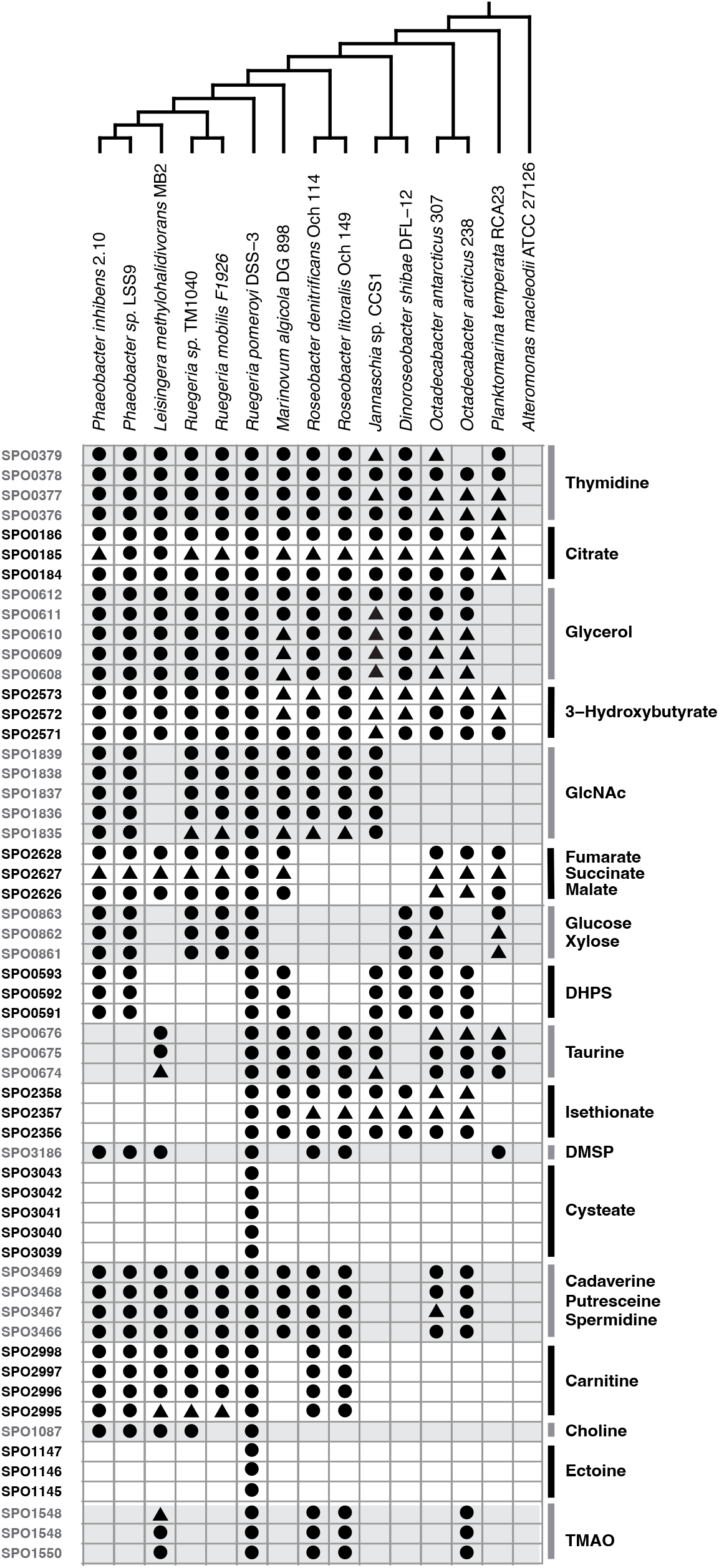
Orthologs of the verified *R. pomeroyi* DSS-3 transporter systems in Roseobacter group members. Each row indicates a single gene and shading indicates genes that make up multi-component transporters. The circles denote orthologs identified by BLASTp using e ≤ 10^−5^ and identity ≥70% thresholds. The triangles denote orthologs of multicomponent that did not meet the BLAST thresholds but were co-located in a transporter operon with components that did. Strain phylogeny is based on analysis of 117 single copy genes.

## Conclusions

Thirteen *R. pomeroyi* transporter annotations were confirmed in a screen of 156 transporter gene mutants (disrupting 104 of the bacterium’s 126 organic carbon influx transporter systems) against 63 metabolites. The verified gene functions provided new insights into in a longitudinal dataset of *R. pomeroyi* transcription through a natural phytoplankton bloom, revealing details of the metabolite landscape and generating the hypothesis that citrate, 3-hydroxybutyrate, taurine, and DMSP were highly available in the dinoflagellate-dominated Monterey Bay bloom. Comparative analysis of the verified transporters across Roseobacter genomes revealed, on the one hand, narrow niche dimensions restricted to subgroups (e.g., *R. pomeroyi* and its closest relatives), and on the other, broad ecological characteristics common across the group and reflecting its core ecological roles. As is the case for many marine bacterial taxa (73), the streamlined Roseobacter species that are more numerous in ocean microbial communities are poorly represented in culture collections (74). As such, experimental gene annotation is key for analyzing, or re-analyzing, microbial gene, transcript, and protein data harboring extensive untapped knowledge among their unannotated genes. For *R. pomeroyi*, this effort brings the percent of organic compound influx transporters with identified substrates to 13% of the 126 gene systems able to acquire metabolites from the ocean’s carbon pools (Table S1).

## Supporting information

Supplemental Figures, merged

## Acknowledgements

The authors thank C. Smith, C. Sanlatte, and J. Schreier for advice and assistance. This work was supported by Simons Foundation grant 542391 to MAM within the Principles of Microbial Ecosystems Collaborative, and NSF award OCE-2019589 to MAM. This is the Center for Chemical Currencies of a Microbial Planet (C-CoMP) publication #00X.

## Data Availability

All growth and BarSeq data are available through BCO-DMO project 884792. All raw NMR data, processing scripts, and processed files for the metabolite drawdown experiment are available in Metabolomics Workbench with Study ID ST002381 (DOI: http://dx.doi.org/10.21228/M8ST4T).

## Contributions

WFS, MAM, and CRR conceived of and designed the research; WFS, HEK, CM, and LTR performed the experiments; MU performed NMR analysis; CM, LTR, and CRR generated and arrayed the mutant library; WFS and MAM conducted statistical analysis, generated figures, and wrote the manuscript with constructive input from all co-authors.

