## Supplemental Figures, merged for "Function and Importance of Marine Bacterial Transporters of Plankton Exometabolites"

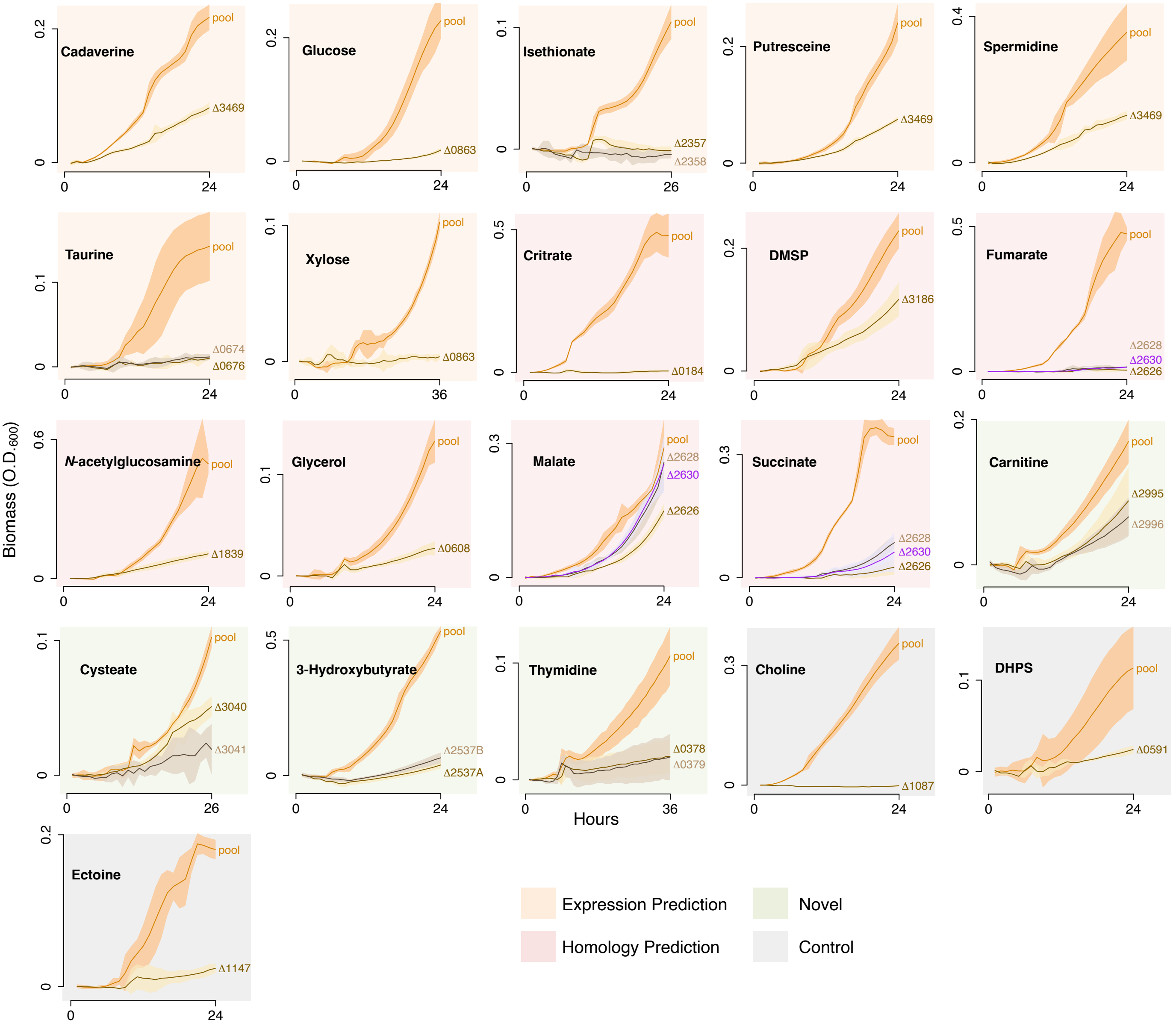


Fig. S1. Growth of individual transporter mutants and the pooled-BarSeq library (an analog for wild-type growth) on selected marine plankton metabolites. Substrates are grouped by previous annotation status of the transporter: orange, predicted by expression; pink, predicted by homology; green, novel; gray, positive controls. Shaded regions indicate 95% confidence intervals (n=4). Numbers refer to *Ruegeria pomeroyi* DSS-3 locus tags (Table 1). Significance indications are from ANOVA and TukeyHSD tests ($p \leq 0.05$) for difference in $OD_{600}$ between pooled-BarSeq and isolated mutants. DMSP, dimethylsulfoniopropionate; DHPS, dihydroxypropanesulfonate.

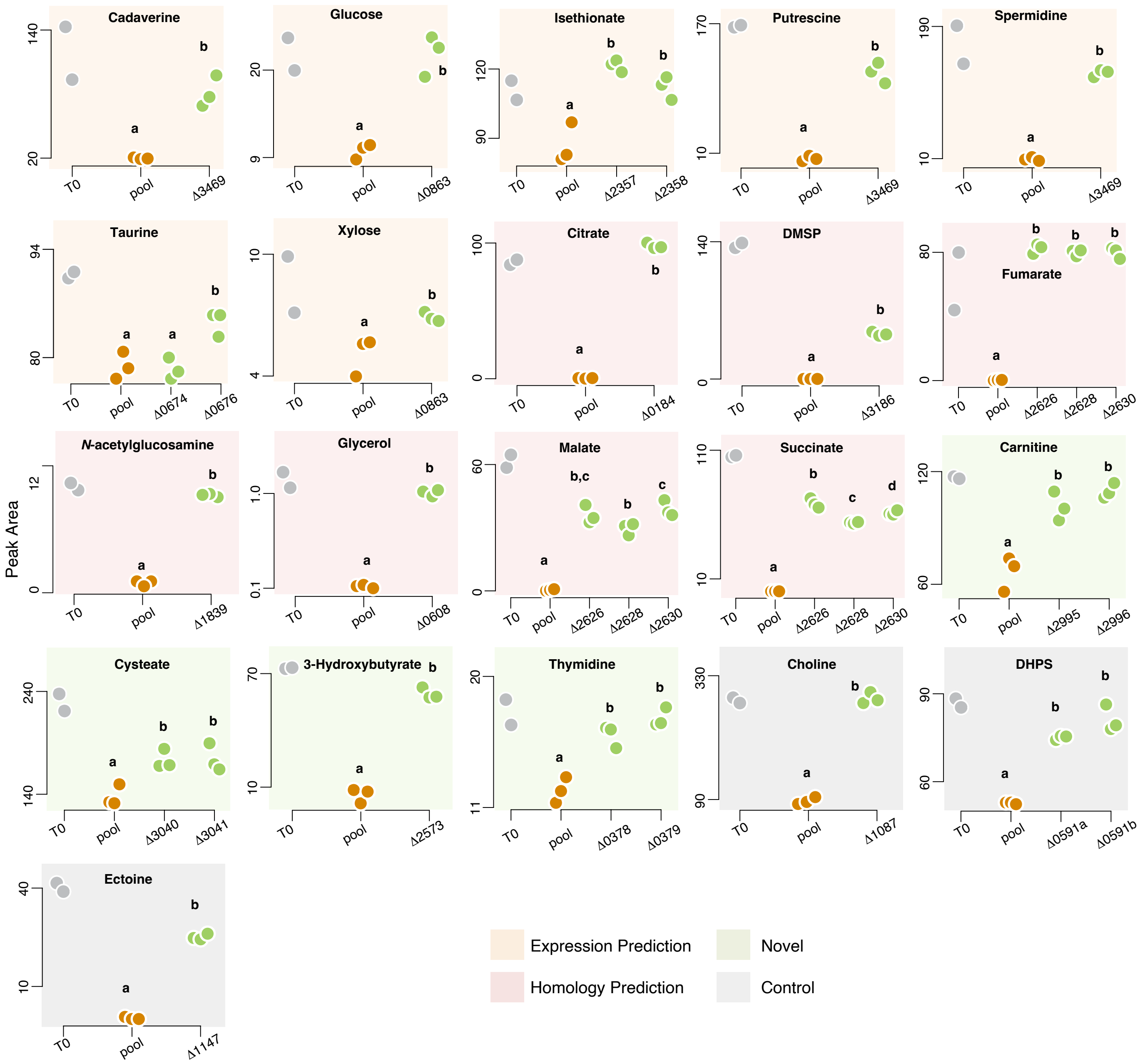


Fig. S2. Substrate concentrations during growth of individual transporter mutants and the pooled-BarSeq library (an analog for wild-type growth) on selected marine plankton metabolites. T0, substrate concentration immediately after inoculation (n=2); pool, substrate concentration after BarSeq mutant pool growth (n=3); Δxxxx, substrate concentration after mutant growth, xxxx indicates *Ruegeria pomeroyi* DSS-3 locus tag (Table 1) (n=3). Plots are grouped by the previous annotation status of the transporter: orange, predicted by expression; pink, predicted by homology; green, novel; gray, positive control. Significance indications are from ANOVA and TukeyHSD tests ($p \leq 0.05$) for difference in peak area between BarSeq and isolated mutants. For full results, see Table 1.

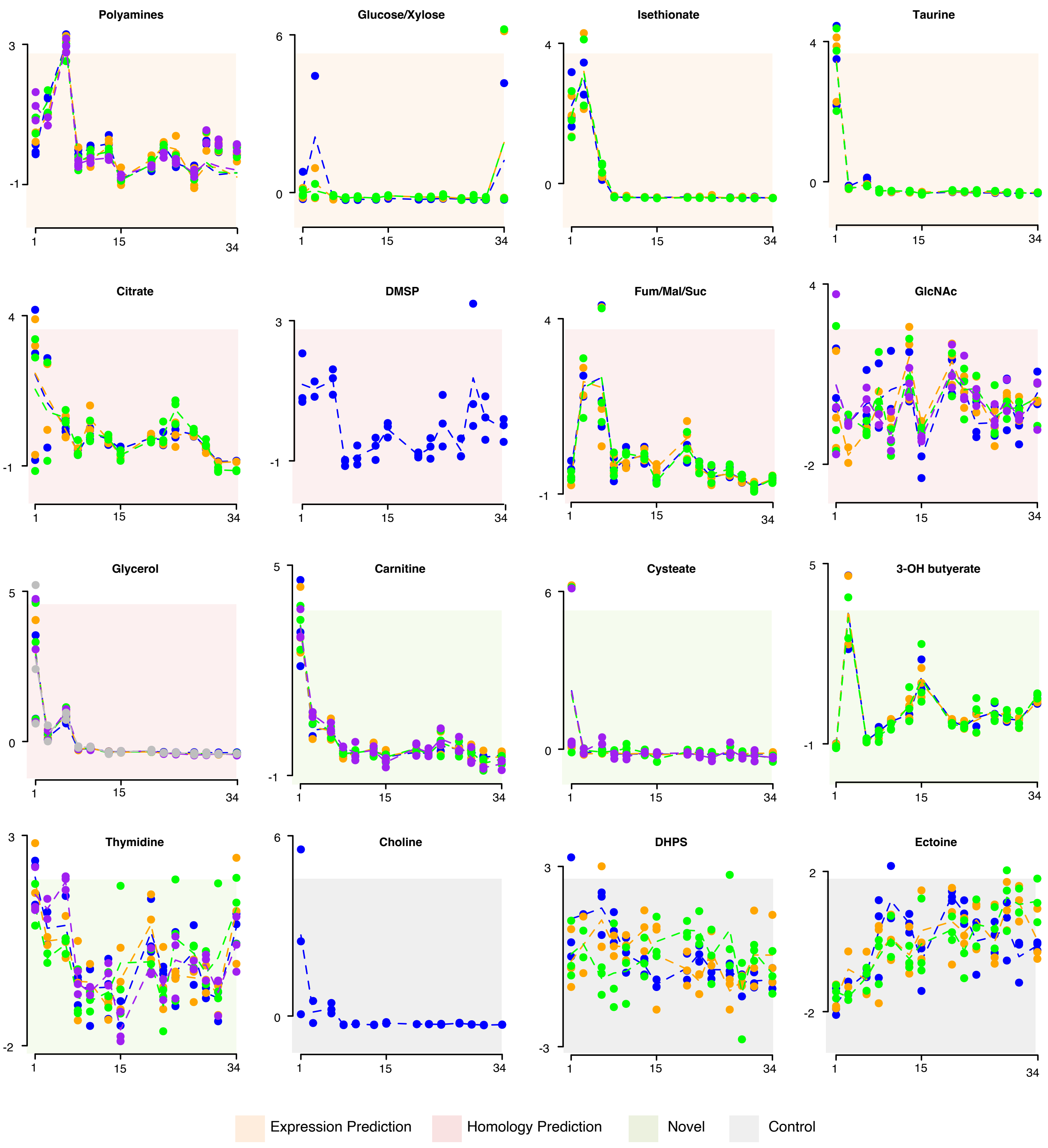


Fig. S3. Relative expression of selected *R. pomeroyi* DSS-3 transporters in Monterey Bay seawater on each of 14 dates over 35 days in Fall 2016, normalized as Z-scores. Transporter component genes (ranging from 1 to 5) with three replicates per component are plotted individually. Lines connect the component mean expression through time.
