## Supplemental Table 1 for "Function and Importance of Marine Bacterial Transporters of Plankton Exometabolites"

**Table S1. Organic compound influx transporter systems in the *Ruegeria pomeryi* DSS-3 genome.**

| Transporter ID | Transporter | Name | Mutant available (at least one component) | Type | Substrate confirmed, this study | Previous annotation (hypothesized or confirmed) | Method for prior annotation | Source |
|---|---|---|---|---|---|---|---|---|
| Transp_001 | SPO0050-0053 ABC transporter | | yes | ABC | | | | |
| Transp_002 | SPO0055 OmpA domain protein | | yes | OmpA | | | | |
| Transp_003 | SPO0065 peptide/opine/nickel uptake family ABC transporter | | | ABC | | | | |
| Transp_004 | SPO0098-0101 peptide/opine/nickel uptake family ABC transporter | | yes | ABC | | | | |
| Transp_005 | SPO0109-0111 ABC-2 type transport system | | yes | ABC | | | | |
| Transp_006 | SPO0123 MFS transporter | | yes | MFS | | | | |
| Transp_007 | SPO0184-00186 tripartate tricarboxylate transporter | *tctABC* | yes | TTT | Citrate | Citrate | GapMind | |
| Transp_008 | SPO0237-0240 glycerol-3-phosphate ABC transporter | | yes | ABC | | | | |
| Transp_009 | SPO0376-0379 sugar or nucleotide ABC transporter | *nupABC* | yes | ABC | Thymidine | novel | | |
| Transp_010 | SPO0398 amino acid/amide ABC transporter | | | ABC | | | | |
| Transp_011 | SPO0472 putative phosphonate transport system | | yes | ABC | | | | |
| Transp_012 | SPO0519-0522 glutamate/glutamine/aspartate/asparagine ABC transporter | | yes | ABC | | | | |
| Transp_013 | SPO0558-0560 oligopeptide ABC transporter | | yes | ABC | | | | |
| Transp_014 | SPO0591-0593 Dihydroxypropanesulfonate (DHPS) TRAP transporter | *hspKLM* | yes | TRAP | DHPS | DHPS | Expression, K | Mayer et al. 2010, Landa et al. 2017 |
| Transp_015 | SPO0608-0612 sugar ABC transporter | *glpVSTPQ* | yes | ABC | Glycerol | Glycerol | GapMind | |
| Transp_016 | SPO0648-0651 sugar ABC transporter | | yes | ABC | | | | |
| Transp_017 | SPO0660-0664 N-acetyltaurine ABC transporter | | yes | ABC | | N-acetyltaurine | Expression | Denger et atl 2011 |
| Transp_018 | SPO0674-0676 taurine ABC transporter | *tauABC* | yes | ABC | Taurine | Taurine | Expression | Gorzynska et al 2006 |
| Transp_019 | SPO0702-0706 oligopeptide ABC transporter | | yes | ABC | | | | |
| Transp_020 | SPO0714-0715 PTS system IIA component, Man family | | | PTS | | | | |
| Transp_021 | SPO0736-0738 TRAP dicarboxylate transporter | | yes | TRAP | | | | |
| Transp_022 | SPO0769-0771 Tripartite-type tricarboxylate transporter | | yes | TTT | | | | |
| Transp_023 | SPO0780-0783 phosphonate ABC transporter | | | ABC | | | | |
| Transp_024 | SPO0789 Hemin uptake protein HemP | | | HemP | | | | |
| Transp_025 | SPO0822-0825 branched-chain amino acid ABC transporter | | yes | ABC | | | | |
| Transp_026 | SPO0861-0863 xylose ABC transporter | *xylFGH* | yes | ABC | Glucose/xylos | Glucose/xylose | Homology, E› | Wiegmann et al 2014 |
| Transp_027 | SPO0874 xanthine/uracil permease | | | xanthine/uracil permease family | | | | |
| Transp_028 | SPO0959 DMT transporter | | yes | DMT | | | | |
| Transp_029 | SPO0973 LAO/AO transport system ATPase | | yes | AAA | | | | |
| Transp_030 | SPO1017-1021 branched-chain amino acid ABC transporter | | yes | ABC | | | | |
| Transp_031 | SPO1028-1029 YeeE/YedE family protein | | | YeeE/YedE | | | | |
| Transp_032 | SPO1060-1062 amino acid ABC transporter | | yes | ABC | | | | |
| Transp_033 | SPO1073-1075 Tripartite-type tricarboxylate transporter | | yes | TTT | | | | |
| Transp_034 | SPO1076 SLC13 family permease | | Yes | SLC13 family | | | | |
| Transp_035 | SPO1087 Choline transporter | *betT* | Yes | BCCT | Choline | Choline | Knockout | Lidbury et al 2015 |
| Transp_036 | SPO1112-1114 C4 dicarboxylate TRAP transporter | | yes | TRAP | | | | |
| Transp_037 | SPO1131-1133 glycine betaine/proline ABC transporter | | yes | ABC | | | | |
| Transp_038 | SPO1145-1147 Ectoine/5-hydroxyectoine TRAP transporter | *uehABC* | yes | TRAP | Ectoine | Ectoine | Knockout | Schultz et al 2016 |
| Transp_039 | SPO1197 amino acid/polyamine/organocation transporter | | yes | APC superfamily | | | | |
| Transp_040 | SPO1210-1213 oligopeptide ABC transporter | | yes | ABC | | | | |
| Transp_041 | SPO1304-1307 His/Glu/Gln/Arg/opine family ABC transporter | | yes | ABC | | | | |
| Transp_042 | SPO1308 ABC transporter; phosphonate | | | ABC | | | | |
| Transp_043 | SPO1404 major facilitator superfamily (MFS) of membrane transport proteins | | yes | MFS | | | | |
| Transp_044 | SPO1454-1456 TRAP C4 dicarboxylate transporter | | yes | TRAP | | | | |
| Transp_045 | SPO1463-1465 TRAP dicarboxylate transporter | | yes | TRAP | | | | |
| Transp_046 | SPO1490-1493 branched-chain amino acid ABC transporter | | yes | ABC | | | | |
| Transp_047 | SPO1514 amino acid/amide ABC transporter | | Yes | ABC | | | | |
| Transp_048 | SPO1516 amino acid ABC transporter | | | ABC | | | | |
| Transp_049 | SPO1543-1547 peptide/opine/nickel uptake family ABC transporter | | yes | ABC | | | | |
| Transp_050 | SPO1548-1550 TMAO ABC transporter | *tmoXYV* | yes | ABC | | TMAO | Knockout | Lidbury et al 2014 |
| Transp_051 | SPO1552 ABC transporter | | yes | ABC | | | | |
| Transp_052 | SPO1606-1609 spermidine/putrescine ABC transporter | | yes | ABC | | | | |
| Transp_053 | SPO1644-1647 oligopeptide/dipeptide ABC transporter | | yes | ABC | | | | |
| Transp_054 | SPO1656-1659 oligopeptide/dipeptide ABC transporter | | yes | ABC | | | | |
| Transp_055 | SPO1707-1710 branched-chain amino acid ABC transporter | | yes | ABC | | | | |
| Transp_056 | SPO1719-1721 TRAP dicarboxylate transporter | | yes | TRAP | | | | |
| Transp_057 | SPO1771-1773 TRAP dicarboxylate transporter | | yes | TRAP | | | | |
| Transp_058 | SPO1785-1788 ABC transporter, taurine or sulfonate-like | | yes | ABC | | | | |
| Transp_059 | SPO1810 sodium/solute symporter, acetate | | | sodium/solute symporter | | | | |
| Transp_060 | SPO1814-1816 TRAP C4 dicarboxylate transporter | | | TRAP | | | | |
| Transp_061 | SPO1820-1823 carbohydrate ABC transporter | | yes | ABC | | | | |
| Transp_062 | SPO1829-1833 branched-chain amino acid ABC transporter | | yes | ABC | | | | |
| Transp_063 | SPO1835-1839 carbohydrate ABC transporter | *nagTUVW* | yes | ABC | GlcNac | GlcNac | GapMind | |
| Transp_064 | SPO1846-1851 branched-chain amino acid ABC transporter | | yes | ABC | | | | |
| Transp_065 | SPO1935-1939 branched-chain amino acid ABC transporter | | yes | ABC | | | | |
| Transp_066 | SPO2006-2009 spermidine/putrescine ABC transporter | | yes | ABC | | | | |
| Transp_067 | SPO2066 ABC peptide/nickel transporter | | | ABC | | | | |
| Transp_068 | SPO2111-2112 OMPP1/FadL/TodX family/DMT | | yes | DMT | | | | |
| Transp_069 | SPO2155 competence protein | | yes | ComEC | | | | |
| Transp_070 | SPO2186-2187 TRAP transporter | | yes | TRAP | | | | |
| Transp_071 | SPO2302 biotin transporter bioY family protein | | | bioY | | | | |
| Transp_072 | SPO2356-2358 Isethionate TRAP transporter | *iseKLM* | yes | TRAP | Isethionate | Isethionate | Expression | Weinitsche et al 2010 |
| Transp_073 | SPO2364-2367 amino acid ABC transporter | | yes | ABC | | | | |
| Transp_074 | SPO2382-2384 tricarboxylate transporter family protein | | yes | TTT | | | | |
| Transp_075 | SPO2431-2433 TRAP dicarboxylate transporter | | yes | TRAP | | | | |
| Transp_076 | SPO2441-2443 glycine betaine/proline ABC transporter | | yes | ABC | | | | |
| Transp_077 | SPO2530-2534 branched-chain amino acid ABC transporter | | yes | ABC | | | | |
| Transp_078 | SPO2545-2547 TRAP dicarboxylate transporter | | yes | TRAP | | | | |
| Transp_079 | SPO2551-2554 peptide/opine/nickel uptake family ABC transporter | | yes | ABC | | | | |
| Transp_080 | SPO2571-2573 TRAP 3-OH butryate | *hbtABC* | yes | TRAP | 3-Hydroxybut‹ | novel | | |
| Transp_081 | SPO2604-2606 hypothetical protein | | yes | TRAP | | | | |
| Transp_082 | SPO2626-2628 TRAP transporter | *dctMPQ* | yes | TRAP | Fumarate, Ma | Fumarate, Mala‹ | GapMind | |
| Transp_083 | SPO2657-2661 cysteate ABC transporter | | yes | ABC | | | | |
| Transp_084 | SPO2664-2667 polar amino acid uptake family ABC transporter | | yes | ABC | | | | |
| Transp_085 | SPO2699-2702 opine/polyamine ABC transporter | | yes | ABC | | | | |
| Transp_086 | SPO2744 malonate transporter, putative | | yes | AEC | | | | |
| Transp_087 | SPO2802-2805 bmp family protein | | yes | BMP | | | | |
| Transp_088 | SPO2813-2816 peptide/nickel/opine uptake family ABC transporter | | yes | ABC | | | | |
| Transp_089 | SPO2831-2835 oligopeptide ABC transporter | | yes | ABC | | | | |
| Transp_090 | SPO2952 trkA domain protein | | yes | trkA | | | | |
| Transp_091 | SPO2995-2998 peptide/nickel/opine uptake family ABC transporter | *cntTUVWX* | yes | ABC | Carnitine | novel | | |
| Transp_092 | SPO3040-3043 polar amino acid uptake family ABC transporter | *cuyTUVW* | yes | ABC | Cysteate | novel | | |
| Transp_093 | SPO3046-3049 oligopeptide ABC transporter | | yes | ABC | | | | |
| Transp_094 | SPO3186 glycine betaine transporter | *dmdT* | yes | BCCT | DMSP | DMSP | Homology | |
| Transp_095 | SPO3290-3295* (not 3293) branched-chain amino acid ABC transporte | | yes | ABC | | | | |
| Transp_096 | SPO3335 glutamine ABC transporter | | yes | ABC | | | | |
| Transp_097 | SPO3339 bioY family protein | | yes | bioY | | | | |
| Transp_098 | SPO3378 benzoate transporter | | | BenE | | | | |
| Transp_099 | SPO3466-3469 putrescine ABC transporter | *potFGHI* | yes | ABC | Spermidine, c | spermidine | Expression | Mou et al 2010 |
| Transp_100 | SPO3472-3475 polyamine ABC transporter | | yes | ABC | | | | |
| Transp_101 | SPO3534-3537 oligopeptide/dipeptide uptake family ABC transporter | | | ABC | | | | |
| Transp_102 | SPO3618-3620 sulfonate ABC transporter | | yes | ABC | | | | |
| Transp_103 | SPO3643 DMT | | | DMT | | | | |
| Transp_104 | SPO3662-3665 hypothetical protein | | yes | TRAP | | | | |
| Transp_105 | SPO3693-3695 TRAP dicarboxylate transporter | | yes | TRAP | | | | |
| Transp_106 | SPO3705-3709 branched-chain amino acid ABC transporter | | yes | ABC | | | | |
| Transp_107 | SPO3774-3778 oligopeptide/dipeptide ABC transporter | | yes | ABC | | | | |
| Transp_108 | SPO3783-3787 sugar ABC transporter | | yes | ABC | | | | |

| | | | |
|---|---|---|---|
| Transp_109 | SPO3843 malonate transporter, putative | | AEC |
| Transp_110 | SPOA0068-0071 polar amino acid ABC transporter | | ABC |
| Transp_111 | SPOA0097-0101 branched-chain amino acid ABC transporter | | ABC |
| Transp_112 | SPOA0118-0120 hypothetical protein | | TTT |
| Transp_113 | SPOA0160-0162 TRAP dicarboxylate transporter | | TRAP |
| Transp_114 | SPOA0166-0167 TRAP transporter solute receptor TAXI family protein | yes | TRAP |
| Transp_115 | SPOA0231-0233 glycine betaine/proline ABC transporter | yes | ABC |
| Transp_116 | SPOA0238-0240 TRAP dicarboxylate transporter | | TRAP |
| Transp_117 | SPOA0249-0251 TRAP dicarboxylate transporter | yes | TRAP |
| Transp_118 | SPOA0253-0258 ribose ABC transporter | yes | ABC |
| Transp_119 | SPOA0263-0265 TRAP transporter | yes | TRAP |
| Transp_120 | SPOA0278-0280 TRAP dicarboxylate transporter | yes | TRAP |
| Transp_121 | SPOA0296-0300 branched-chain amino acid ABC transporter | yes | ABC |
| Transp_122 | SPOA0333-0335 TRAP dicarboxylate transporter | yes | TRAP |
| Transp_123 | SPOA0349 DMT family transporter | yes | DMT |
| Transp_124 | SPOA0366-03667 amino acid permease | yes | APC |
| Transp_125 | SPOA0372-0374 TRAP dicarboxylate transporter | yes | TRAP |
| Transp_126 | SPOA0381-0384 spermidine/putrescine ABC transporter | yes | ABC |
